# Variable impact of *Verticillium longisporum* on oilseed rape yield in field trials in the United Kingdom

**DOI:** 10.1101/205401

**Authors:** Jasper R. L. Depotter, Bart P.H.J. Thomma, Thomas A. Wood

## Abstract

The *Verticillium* genus comprises economically important plant pathogens that collectively affect a broad range of annual and perennial crops. *Verticillium longisporum* mainly infects brassicaceous hosts, including oilseed rape. The most conspicuous symptom of *V. longisporum* infection on oilseed rape is black stem striping that appears towards the end of the cropping season. Thus far, the impact of *V. longisporum* infection on oilseed yield remains unclear. Verticillium stem striping recently emerged as a new disease in UK and currently displays a widespread occurrence in England. In this study, we assessed the impact of Verticillium stem striping on British oilseed rape production. To this end, four cultivars (Incentive, Vision, Harper and Quartz) were grown in field plots with different levels of *V. longisporum* disease pressure at different locations over two consecutive years. Whereas Incentive and Vision developed relatively few stem striping symptoms, Harper and especially Quartz showed severe symptoms during these field experiments. Furthermore, higher inoculum levels induced more severe symptoms in these cultivars. Intriguingly, significant yield reductions upon *V. longisporum* infection only occurred in a single field trial on all tested oilseed rape cultivars. Thus we conclude that Verticillium stem striping does not consistently impact oilseed rape yield, despite the occurrence of abundant disease symptoms.

## Introduction

Oilseed rape is the second most important oilseed crop worldwide with an annual production of over 70 megatons in 2014 (FAOSTAT, 2016). Rape oil is used for many purposes, such as in human consumption, as bio-fuel, and as a protein source for animal feed (Berry et al., 2014). Global oilseed rape production has boomed over the last few decades with the cultivated area increasing over 80% between 1993 and 2013 (FAOSTAT, 2016). A similar trend was observed in the United Kingdom (UK), where the crop developed from being practically unknown before 1970, to the third most widely cultivated crop with an approximate annual production area of 700,000 hectares at present (Wood et al., 2013). Oilseed rape is often grown in increasingly shortening rotation schemes with cereal crops, such as barley (*Hordeum vulgare*) and wheat (*Triticum aestivum*) (Rathke et al., 2005; Wallenhammar et al., 2014). Contemporary oilseed rape cultivars can be divided into two types: open pollinated varieties and hybrid varieties (Wood et al., 2013). Hybrid oilseed cultivars benefit from heterosis and generally have higher, more stable yield than their parents (Diepenbrock, 2000). Commercial hybrid cultivars generally show greater early season vigour and produce higher yields than open pollinating cultivars (Wood et al., 2013).

Oilseed rape production systems are threatened by a plethora of pathogens. Stem canker (*Leptosphaeria spp.*) and light leaf spot (*Pyrenopeziza brassicae*) are considered the most important pathogens for British oilseed rape (Fitt et al., 2006). However, Verticillium stem striping is a newly emerging oilseed rape disease in the UK and its impact on yield is unclear (Dunker et al., 2008; Gladders et al., 2011). Verticillium stem striping is caused by the fungal pathogen *Verticillium longisporum*, one of the ten species of the *Verticillium* genus (Inderbitzin et al., 2011). *Verticillium* spp. comprises several wilt agents on economically important crops, of which *V. dahliae* is the most notorious one because of its extremely wide host range that comprises hundreds of species (Fradin and Thomma, 2006; Inderbitzin and Subbarao, 2014). In contrast to *V. dahliae*, the host range of *V. longisporum* is more confined and mainly comprises brassicaceous plants (Eynck et al., 2007; Inderbitzin and Subbarao, 2014; Novakazi et al., 2015; Zeise and Tiedemann, 2002). On these hosts, *V. longisporum* causes wilting symptoms (Koike et al., 1994). However, no wilting symptoms are observed on oilseed rape (Heale and Karapapa, 1999). Rather, dark unilateral striping is induced at the end of the cropping season before the onset of maturation, approximately 3-4 weeks before harvest. This coincides with the formation of black microsclerotia by the fungus in the stem cortex underneath the epidermis. Hence, the general name “Verticillium stem striping” was coined for this disease on oilseed rape (Depotter et al., 2016). The late appearance of the stem striping symptoms may be related to differences in xylem sap constitution during oilseed rape development, as xylem sap of older plants is more conducive for *V. longisporum* growth (Lopisso et al., 2017a).

The first report of Verticillium stem striping dates from 1969 in the west and south of Scania, southern Sweden, and the disease is currently well established in north-central Europe (Heale and Karapapa, 1999; Kroeker, 1970; Steventon et al., 2002; Zhou et al., 2006). In 2007, Verticillium stem striping was identified for the first time in the UK and is currently present throughout England (Depotter et al. in press b; Gladders et al., 2013, 2011). The cause of this sudden emergence of Verticillium stem striping remains unknown. The British *V. longisporum* population displays a large genetic diversity when compared with populations from other countries, indicating that the presence of *V. longisporum* predates 2007 (Depotter et al. in press b). In support of this, *V. longisporum* was previously found during the 1950s, albeit on a different host: Brussels sprout (*Brassica oleracea* var. *gemmifera*) (Isaac, 1957). Thus, the sudden rise of Verticillium stem striping may perhaps be caused by altered environmental factors in combination with an increase in the area of oilseed rape cultivated.

The impact of Verticillium stem striping on crop quality and quantity is under-investigated (Dunker et al., 2008). Yield losses by Verticillium stem striping have been anticipated to range from 10 to 50%, yet experimental verifications of such estimations are lacking (Dunker et al., 2008). In contrast, Verticillium stem striping did not impact yield significantly in previous field studies despite the presence of stem striping symptoms (Dunker et al., 2008). Nevertheless, pathogenicity tests under controlled conditions demonstrate the destructive potential of *V. longisporum* on oilseed rape (Depotter et al. in press a; Novakazi et al., 2015).

In this study, we assessed the impact of Verticillium stem striping on British oilseed rape production in field experiments. Symptom development was monitored on four different oilseed rape cultivars and yield data were recorded in order to investigate putative differences in cultivar responsiveness to *V. longisporum*.

## Materials and methods

### Field experiments

Field experiments were performed in two consecutive years at different locations. In harvest year 2016, field plots were located in Hinxton, Cambridgeshire, UK, whereas in harvest year 2017 fields were situated in Higham, Suffolk, UK. Four winter oilseed rape cultivars were tested of which two were open-pollinated: Vision and Quartz, and two were hybrid types: Incentive and Harper. The experiments were arranged in a semi-randomized block design with four replicate plots of 24 m^2^ in size for every treatment and cultivar, totalling 48 plots. Buffer zones were included in the experimental design between plots with different inoculum levels to prevent contamination of *V. longisporum* propagules to adjacent plots. The test plots were inoculated with different quantities of *V. longisporum* microsclerotia. To this end, a mix of British *Verticillium* strains were cultured in bags containing a sterile, moist medium of vermiculite (800 ml) and maize meal (26 g) for approximately four weeks and then dried at room temperature for a week prior to inoculation. The plots were either not inoculated, or inoculated by homogenously spreading 0.25 l or 2.5 l of thoroughly mixed, colonized vermiculite/maize meal medium. To enable homogenous dispersion, the 0.25 l of inoculum was first mixed with 0.75 l sand. Inoculations were performed the 16^th^ of September 2015 for the field trial at Hinxton and the 26^th^ of September 2016 for the trial at Higham. The field plots were sown 5^th^ and 21^st^ September, and harvested on 31^th^ and 26^th^ July for the field trials at Hinxton and Higham, respectively. To this end, a central strip of 18 m^2^ was harvested using a combine and used for analysis. Seed yield of oilseed rape were adjusted to 9% moisture content and oil content of the seeds was determined by Nuclear Magnetic Resonance (NMR) spectroscopy (Benchtop NMR Analyser – MQC, Oxford Instruments, UK).

### Disease scoring

Disease scores were based on the fraction of stem circumference that displayed Verticillium stem striping symptoms. For every plot at each time point, 20 stems were randomly chosen and scored on a 0-10 scale, with each interval representing a 10% increase in dark unilateral striping of the stem circumference (eg. Score 2 = 20% striping). The symptoms were scored four times each year starting from the end of June with a 7-day interval: scoring point 1 on 30^th^ June and 27^th^ June, point 2 on 7^th^ July and 4^th^ July, point 3 on 14^th^ July and 11^th^ July, and point 4 on 21^st^ July and 18^th^ July for year 2016 and 2017, respectively.

### Data analysis

Data analysis was performed with R3.2.3 (R Core Team, 2015). Significance levels and correlations were calculated parametrically through the one-way analysis of variance (ANOVA) and Pearson’s correlation coefficient (*r*), respectively. Non-parametric differences in significance were determined through Mann-Whitney U-test or with bootstrap replications of median differences. Correlations of non-parametric data were calculated with the Spearman’s correlation coefficient (*ρ*). Area under disease progress curve (AUDPC) was calculated by trapezoidal integration to compare disease severities over time between different years, cultivars and disease levels (Shaner and Finney, 1977).

## Results

The impact of Verticillium stem striping disease in oilseed rape fields was assessed on four different cultivars: the hybrid cultivars Incentive and Vision, and the open pollinating cultivars Harper and Quartz, by recording stem striping symptoms. During the cropping season clear differences in cultivar responses were observed (Figure 1). The cultivars Incentive and Vision displayed equally few disease symptoms at the end of both cropping seasons (Table 1). Consequently, little to no progress in disease symptoms was monitored in both years (Figure 1). Despite the low degree of symptom development, significant differences in disease incidence and symptom severity were recorded for the different inoculum amounts (Table 1). Disease symptoms were generally more severe in plots with the higher inoculum treatment when compared with control plots or plots with the lower inoculum amount.

**Figure 1.**
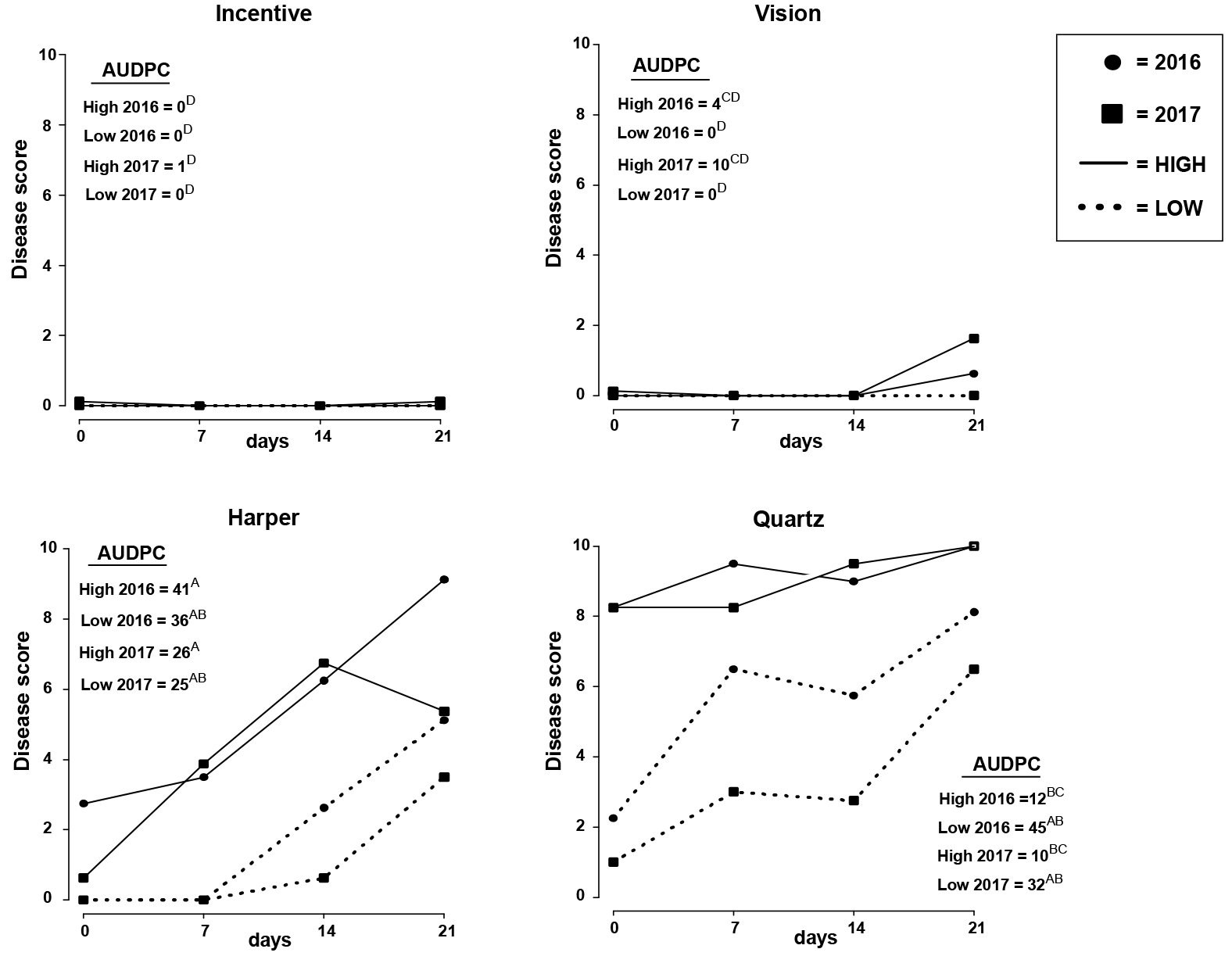
Verticillium stem striping progression during the end of the cropping season. The Y-axis represents the median of the upper quartile plot disease score. The X-axis represents the number of days after the first scoring: 30^th^ June and 27^th^ June for harvest years 2016 and 2017, respectively. The area under progress disease curve (AUPDC) was calculated and letters present significance levels calculated with 50,000 bootstrap replications of median differences (*P*<0.05).

**Table 1.**
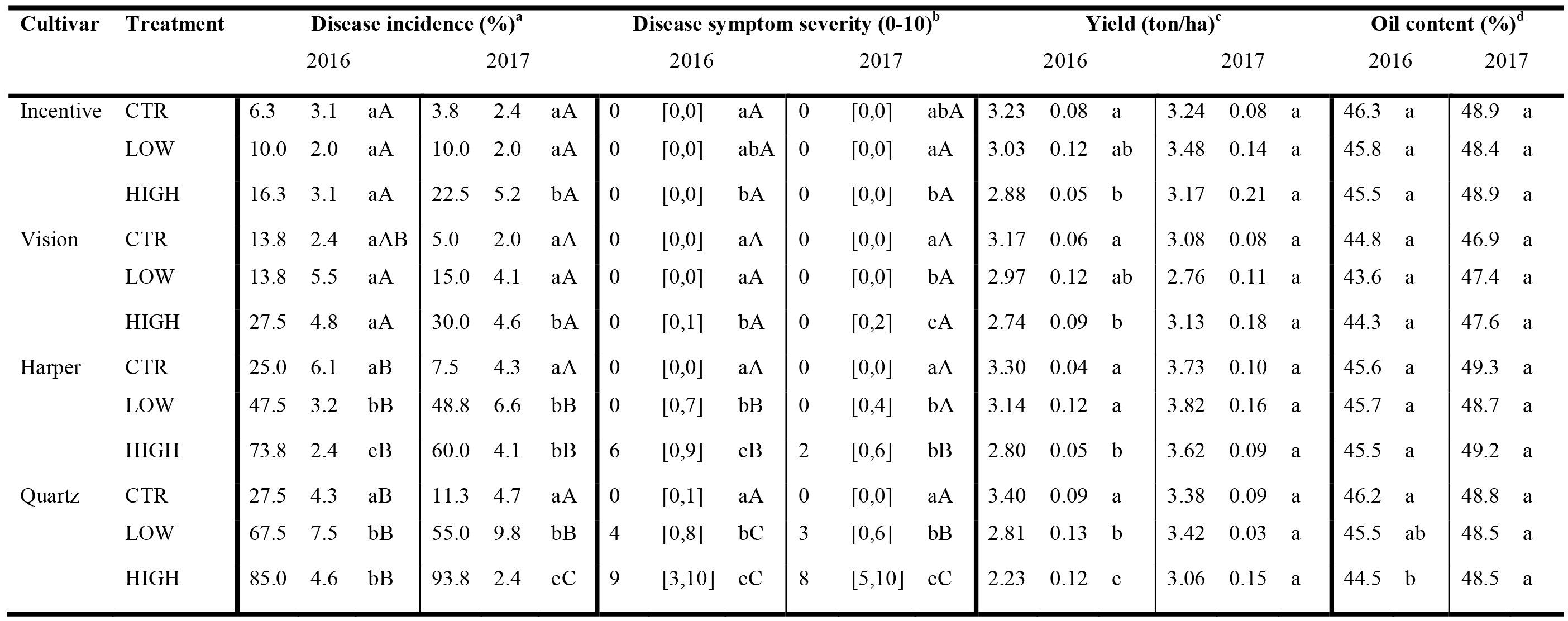
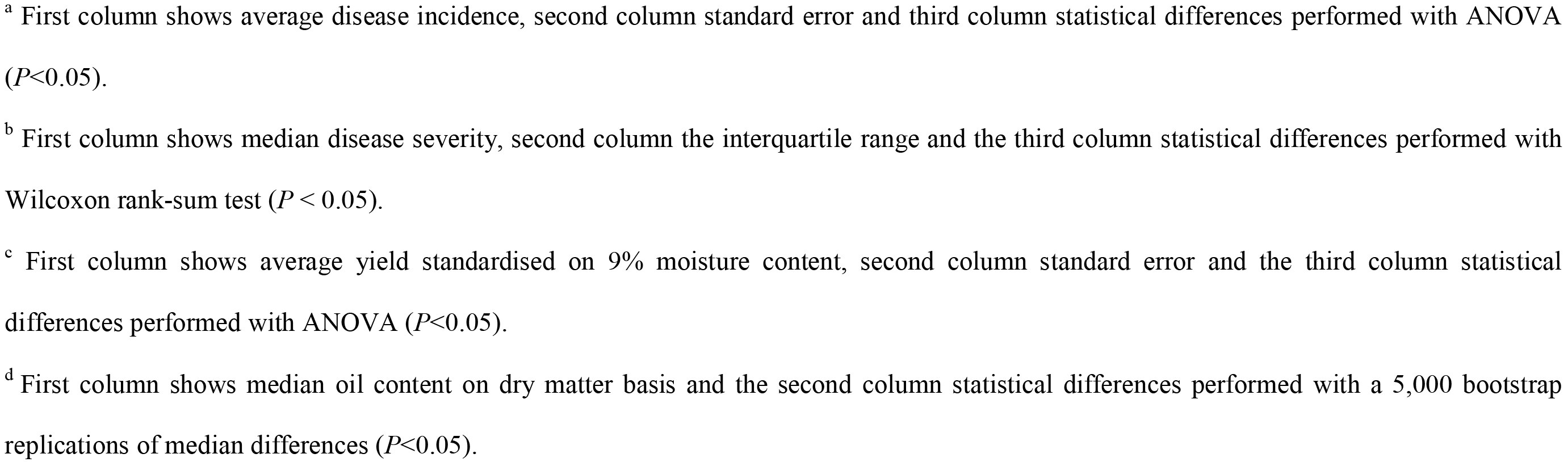
Impact of Verticillium stem striping on oilseed rape: disease symptoms, yield and oil content. Data of disease incidence and symptom severity were assessed on 21 July and 18 July for year 2016 and 2017, respectively. Lower case letters indicate significant differences between treatments of the same cultivar in the same year, whereas capital letters indicate significant differences between cultivars of the same treatment in the same year. The treatment abbreviations “CTR”, “LOW” and “HIGH”, represent the data for plots were no, lower and higher amounts of *V. longisporum* microsclerotia were applied, respectively.

In comparison to the cultivars Incentive and Vision, Harper displayed more extensive disease symptoms in both consecutive trial years (Table 1). Altogether, disease symptom development was similar in both cropping years (Figure 1). Higher inoculum levels resulted in increasing incidence and symptom severity in Harper for the 2016 trial (Table 1). However, whereas disease levels the inoculated plots were higher than in control plots, symptoms levels between the lower and higher inoculum treatments could not be distinguished in 2017 (Table 1).

Of all oilseed rape cultivars tested, Quartz displayed the most severe disease symptoms upon *V. longisporum* challenge (Table 1). The area under disease progress curves (AUDPCs) for the lower and higher inoculum treatments were similar for both years. Although it seems that AUDPCs were higher in plots with lower when compared with higher inoculum levels, and disease symptoms were already severe at the start of monitoring in plots with higher inoculum levels, no statistical methodology could be applied to account for this observation (Figure 1). Higher inoculum levels resulted in increasing incidence and symptom severity in Quartz for both cropping years (Table 1).

In order to evaluate whether differential symptomatology corresponded with economically important crop parameters, seed yield and oil content were measured. Differences in oilseed rape cultivar and trial year impacted significantly on seed yield and oil content, whereas differences in the level of inoculum only impacted on yield and not oil content (Table 2). However, the impact of inoculum level was inconsistent between the two cropping years as significant differences in yield and oil content could only be found in the 2016 field trial (Table 1). In 2016, plots with the higher inoculum rate yielded significantly less than control plots for all tested oilseed rape cultivars. The highest yield impact was for Quartz that demonstrated a 34% reduction compared to the un-inoculated control (Table 1). Despite the presence of only mild disease symptoms, the yield in 2016 was also significantly lower for Incentive (-11%) and Vision (-14%). Intriguingly, the degree of yield reduction was similar for Harper (-15%), despite the observation that disease incidence and severity are significantly higher on this cultivar (Table 1). Yield for plots inoculated with the lower amount of inoculum only a showed significantly reduction for Quartz (-17%) while the other cultivars did not suffer from measurable yield losses (Table 1). The higher yield reductions in Quartz when compared to other cultivars correspond to its higher level of stem striping symptoms. Across all cultivars, the median disease symptom severity negatively correlated to yield (*ρ* = −0.53, *P* = 0.0001). Similarly, the average disease incidence also correlated negatively with yield (*r* = −0.57, *P* = 0).

**Table 2.** Analysis of variance for effects on seed yield and oil content.

|  | Yield |  |  |  |  | Oil content |  |  |  |  |
| --- | --- | --- | --- | --- | --- | --- | --- | --- | --- | --- |
|  | Df | Sum Sq | Mean Sq | F-value | P-value | Df | Sum Sq | Mean Sq | F-value | P-value |
| Main effects |  |  |  |  |  |  |  |  |  |  |
| A: Cultivar | 3 | 2.53 | 0.84 | 20.71 | 4.58e<sup>-6</sup> | 3 | 33.81 | 11.27 | 14.82 | 4.17e<sup>-5</sup> |
| B: Year | 1 | 2.93 | 2.93 | 71.98 | 1.05e<sup>-7</sup> | 1 | 253.27 | 253.27 | 332.91 | 4.67e<sup>-13</sup> |
| C: Inoculum level | 2 | 2.14 | 1.07 | 26.32 | 4.53e<sup>-6</sup> | 2 | 1.19 | 0.60 | 0.79 | 0.47 |
| D: Repeat | 3 | 0.38 | 0.13 | 3.14 | 0.05 | 3 | 9.59 | 3.19 | 4.19 | 0.02 |
| Interactions |  |  |  |  |  |  |  |  |  |  |
| A:B | 3 | 1.27 | 0.42 | 10.40 | 3.4e<sup>-4</sup> | 3 | 0.98 | 0.33 | 0.43 | 0.74 |
| A:C | 6 | 1.11 | 0.18 | 4.54 | 5.67e<sup>-3</sup> | 6 | 3.76 | 0.63 | 0.82 | 0.57 |
| A:D | 9 | 0.20 | 0.02 | 0.53 | 0.83 | 9 | 6.40 | 0.71 | 0.94 | 0.52 |
| B:C | 2 | 1.02 | 0.51 | 12.49 | 3.97e<sup>-4</sup> | 2 | 2.54 | 1.27 | 1.67 | 0.22 |
| B:D | 3 | 0.29 | 0.10 | 2.36 | 0.11 | 3 | 11.04 | 3.68 | 4.84 | 0.01 |
| C:D | 6 | 0.64 | 0.11 | 2.63 | 0.05 | 6 | 3.55 | 0.59 | 0.78 | 0.60 |
| A:B:C | 6 | 0.52 | 0.09 | 0.09 | 0.10 | 6 | 2.93 | 0.49 | 0.64 | 0.70 |
| A:B:D | 9 | 0.15 | 0.02 | 0.41 | 0.91 | 9 | 2.78 | 0.31 | 0.41 | 0.92 |
| A:C:D | 18 | 0.97 | 0.05 | 1.32 | 0.28 | 18 | 8.35 | 0.46 | 0.61 | 0.85 |
| *B:C:D* | 6 | 0.23 | 0.04 | 0.95 | 0.48 | 6 | 3.77 | 0.63 | 0.83 | 0.57 |
| Residual | 18 | 0.73 | 0.04 |  |  | 18 | 13.69 | 0.76 |  |  |
| Total | 95 |  |  |  |  | 95 |  |  |  |  |

In contrast to the observations in 2016, the relationship between Verticillium stem striping symptoms and yield was different in the 2017 trial, since no significant differences between the different inoculum treatments were observed in yield for any of the cultivars assessed (Table 1). Verticillium stem striping symptoms, incidence (*r* = 0.05, *P* = 0.73) and severity (*ρ* = 0.09, *P* = 0.52) did not correlate with yield.

## Discussion

Verticillium stem striping is an elusive disease as its symptoms often remain unnoticed and its impact on oilseed rape production is under-investigated (Depotter et al., 2016). Moreover, its economic significance has been questioned as yield reductions were assumed rather than experimentally verified (Dunker et al., 2008). In this study, we demonstrated the yield-reducing potential of Verticillium stem striping disease under field conditions, as significant yield reductions occurred in the 2016 trial. In 2016, the tested cultivars displayed yield reduction between 11 and 34%, which is within the range of previously made estimates of 10 to 50% (Dunker et al., 2008). Yield correlated negatively with the incidence and severity of stem striping symptoms. In contrast, no impact on yield was observed in the 2017 trial, not even for the Quartz cultivar that developed severe disease symptoms in that year. Moreover, the symptom development on Quartz in 2016 was comparable to that in 2017. This observation suggests that environmental factors impact the occurrence of yield reductions. Weather data was tentatively analysed, as the data were not collected on site but of a weather station located within a 35 km radius with both field locations (Met Office, 2017). No noteworthy differences in temperature and precipitation were observed from October through to February, a period that is important for oilseed rape leaf development. However, during March and April, the average temperature was almost three degrees lower in 2016 (11.2°C) when compared with 2017 (14.0°C). Furthermore, rainfall was over twice as much during March and April in 2016 (109 mm) when compared with 2017 (46 mm), which may have implications on stem extension, flower bud development and flowering onset. In contrast, the final months of the cropping season (May, June and July) were relatively dry in 2016 (124 mm precipitation) when compared with 2017 (213 mm), which may impact pod and seed development, and plant senescence. Little is known of the impact of weather conditions on Verticillium stem striping, and further research is needed to reveal putative relations on the yield impact of this disease. However, watering regimes were previously shown not to impact *V. longisporum* disease levels if grown under controlled greenhouse conditions (Lopisso et al., 2017b).

Verticillium are difficult diseases to control as no curative fungicides are available (Depotter et al., 2016). Hence, apart from alternative management strategies such as crop rotation (Bhat and Subbarao, 1999) and bio-control (Deketelaere et al., 2017), improving cultivar resistance is the most effective way to combat *Verticillium* diseases. This is reflected in the research efforts to improve Verticillium stem striping resistance in oilseed rape (Eynck et al., 2009b; Happstadius et al., 2003; Obermeier et al., 2013; Rygulla et al., 2008, 2007a, 2007b). Cultivar resistance or partial resistance are useful resources in breeding programs to mitigate the effects of disease. Thus, the availability of mildly susceptible cultivars such as Incentive and Vision in comparison to Harper and Quartz is promising for reducing the effects of *V. longisporum* in oilseed crops. A similar difference in susceptibility was previously observed in pathogenicity tests under controlled greenhouse conditions, where Quartz displayed more severe disease symptom development upon *V. longisporum* inoculation when compared with Incentive (Depotter et al. in press a). The understanding of mechanisms responsible for differences in Verticillium stem striping susceptibility in oilseed rape cultivars is still limited. However, a significant role for physical cell wall barriers is anticipated (Eynck et al., 2009a; Lopisso et al., 2017a). Reinforcements of tracheary elements through wall-bound depositions of phenolics and lignin is associated with increased *V. longisporum* resistance on oilseed rape lines, possibly hampering fungal colonization of in the xylem tissue (Eynck et al., 2009a).

Although disease symptoms negatively correlated to yield in the 2016 field trial, yield reductions in Harper were comparable to Incentive and Vision, despite differences in the symptoms displayed (Figure 1, Table 1). In addition, the lack of significant yield reductions in the presence of disease symptoms in the 2017 trial indicates that stem striping is a poor indicator for predicting yield losses by *V. longisporum*. Stem striping symptom development may depend on cultivar-specific physiological properties, such as time-point of ripening, as stem striping symptoms develop during this physiological stage (Depotter et al., 2016). Stem striping may be undesirable for its potential role in inoculum build-up. Oilseed cultivars that display more stem striping symptoms may increase the soil inoculum, as stripes are the consequence of melanized microsclerotia that accumulate in the stem cortex. This inoculum can negatively impact brassicaceous crops in the following years, as increased inoculum levels generally led to more disease symptoms and an increased potential of yield reduction.

Environmental factors or cultural practices are deemed to be responsible for the sudden emergence of Verticillium stem striping in British oilseed rape (Depotter et al. in press b). In this study, we illustrate the importance of oilseed rape cultivar choice with respect to the display of striping symptoms and possible yield impact of *V. longisporum*. Possibly, the anticipated latency and sudden emergence of Verticillium stem striping in the UK may be related to changes in genetic background of currently used oilseed rape cultivars. Arguably, Verticillium stem striping would not be reported in fields where Incentive and Vision are grown, whereas, if symptoms are as abundant as for Quartz, stem striping might be noticed and reported as a new disease.

## ACKNOWLEDGEMENTS

The authors would like to thank the Marie Curie Actions program of the European Commission that financially supports the research of J.R.L.D. Work in the laboratory of B.P.H.J.T. is supported by the Research Council Earth and Life Sciences (ALW) of the Netherlands Organization of Scientific Research (NWO).

